# Genome-wide study identifies association between HLA-B*55:01 and penicillin allergy

**DOI:** 10.1101/2020.02.27.967497

**Authors:** Kristi Krebs, Jonas Bovijn, Maarja Lepamets, Jenny C Censin, Tuuli Jürgenson, Dage Särg, Yang Luo, Line Skotte, Frank Geller, Bjarke Feenstra, Wei Wang, Adam Auton, 23andMe Research Team, Soumya Raychaudhuri, Tõnu Esko, Andres Metspalu, Sven Laur, Michael V Holmes, Cecilia M Lindgren, Reedik Mägi, Lili Milani, João Fadista

**Author notes:** These authors contributed equally (Kristi Krebs, Jonas Bovijn, Michael V Holmes, Cecilia M Lindgren, Reedik Mägi, Lili Milani, João Fadista). Corresponding author, Lili Milani, PhD.

## Abstract

**Background:** Hypersensitivity reactions to drugs are often unpredictable and can be life-threatening, underscoring a need for understanding their underlying mechanisms and risk factors. The extent to which germline genetic variation influences the risk of commonly reported drug allergies such as penicillin allergy remains largely unknown.

**Methods:** We extracted data from the electronic health records of 52,000 Estonian and 500,000 UK biobank participants to study the role of genetic variation in the occurrence of penicillin hypersensitivity reactions. We used imputed SNP to HLA typing data from up to 22,554 and 488,377 individuals from the Estonian and UK cohorts, respectively, to further fine-map the human leukocyte antigen (HLA) association and replicated our results in two additional cohorts involving a total of 1.14 million individuals.

**Results:** Genome-wide meta-analysis of penicillin allergy revealed a significant association located in the HLA region on chromosome 6. The signal was further fine-mapped to the HLA-B*55:01 allele (OR 1.47 95% CI 1.37-1.58, P-value 4.63×10^-26^) and confirmed by independent replication in two cohorts. The meta-analysis of all four cohorts in the study revealed a strong association of HLA-B*55:01 allele with penicillin allergy (OR 1.33 95% CI 1.29-1.37, P-value 2.23×10^-72^). *In silico* follow-up suggests a potential effect on T lymphocytes at HLA-B*55:01.

**Conclusion:** We present the first robust evidence for the role of an allele of the major histocompatibility complex (MHC) I gene HLA-B in the occurrence of penicillin allergy.

## MAIN

Adverse drug reactions (ADRs) are common in clinical practice and are associated with high morbidity and mortality. A meta-analysis of prospective studies in the US revealed the incidence of serious ADRs to be 6.7% among hospitalized patients, and the cause of more than 100,000 deaths annually^1^. In Europe, ADRs are responsible for 3.5% of all hospital admissions, with 10.1% of patients experiencing ADRs during hospitalization and 197,000 fatal cases per year^2,3^. In the US, the cost of a single ADR event falls between 1,439 to 13,462 USD^4^.

ADRs are typically divided into two types of reactions. Type A reactions are more predictable and related to the pharmacological action of a drug, whereas type B reactions are idiosyncratic, less predictable, largely dose-independent, and typically driven by hypersensitivity reactions involving the immune system^5^. Although type B reactions are less frequent (<20%) than type A reactions, they tend to be more severe and more often lead to the withdrawal of a drug from the market^6^. One of the most common causes of type B reactions are antibiotics^5^, typically from the beta-lactam class, with the prevalence of penicillin allergy estimated to be as high as 25% in some settings^7,8^. Despite the relative frequency of such reactions, there are very few studies of the genetic determinants of penicillin allergy^9,10^. This underscores the need for a better understanding of the mechanisms and risk factors, including the role of genetic variation, that contribute to these reactions.

The increasing availability of genetic and phenotypic data in large biobanks provides an opportune means for investigating the role of genetic variation in drug-induced hypersensitivity reactions. In the present study, we sought to identify genetic risk factors underlying penicillin-induced hypersensitivity reactions by harnessing data from the Estonian (EstBB) and UK Biobanks (UKBB), with further replication in two large cohorts.

## METHODS

### Study subjects

We studied individual-level genotypic and phenotypic data of 52,000 participants from the Estonian Biobank (EstBB) and 500,000 participants from UK Biobank (UKBB). Both are population-based cohorts, providing a rich variety of phenotypic and health-related information collected for each participant. We extracted information on penicillin allergy by searching the records of the participants for Z88.0 ICD10 code indicating patient-reported allergy status due to penicillin. Information on phenotypic features like age and gender were obtained from the biobank recruitment records. We also extracted likely penicillin allergies in the EstBB from the recruitment questionnaires and free text fields of the electronic health records (EHRs) using a rule-based approach (see **Supplementary methods** for further details).

### Genome-wide study and meta-analysis

The details on genotyping, quality control and imputation are fully described elsewhere for both EstBB^11,12^ and UKBB^13^. In the Estonian biobank, we conducted the penicillin GWAS among 31,760 unrelated individuals of whom 961 were cases with self-reported allergy to J01C beta-lactam drugs and 30,799 undiagnosed controls. In the UKBB, GWAS on penicillin allergy (Z88.0) was performed among 15,690 cases and 342,116 controls. The analyses were adjusted for the first ten PCs of the genotype matrix, as well as for age, sex and array (see **Supplementary methods**). We performed meta-analysis of 19,051,157 markers (MAF>0.1%) based on effect sizes and their standard errors using METAL^14^. Results were visualized with R software (3.3.2)^15^.

### HLA-typing

HLA-typing of the EstBB genotype data was performed at the Broad Institute using the SNP2HLA tool^16^. The imputation was done for genotype data generated on the GSA, and after quality control the four-digit HLA alleles of 22,554 individuals were used for analysis. In UKBB we used four-digit imputed HLA data released by UKBB^13,17^. The imputation process, performed using HLA*IMP:02^18^, is described more fully elsewhere^13^ and in the **Supplementary methods**.

We performed separate additive logistic regression analysis with the called HLA alleles using R *glm* function in EstBB and UKBB including age, sex and 10 PCs as covariates. Meta-analysis of 162 HLA alleles was performed with the GWAMA software tool^19^. A Bonferroni-corrected P-value threshold of 3.09×10^-4^ was applied based on the number of tested alleles: 0.05/162.

For detection of the strongest tagging SNP for the HLA-B*55:01 allele we calculated Pearson correlation coefficients between the HLA-B*55:01 allele and all the SNPs within +/- 50kb of the HLA-B gene region using R (3.3.2)^15^ *cor* function.

### HLA-B*55:01 replication

Replication analysis of the HLA-B*55:01 allele was tested on 87,996 cases and 1,031,087 controls of European ancestry (close relatives removed) from the 23andMe research cohort using a logistic regression assuming an additive model (see details in the **Supplementary methods).**The self-reported phenotype of penicillin allergy was defined as an allergy test or allergic symptoms required for cases, with controls having no allergy. Estimates from BioVu were extracted from the BioVu publicly available data resource (https://phewascatalog.org/hla). Meta-analysis of the HLA-B*55:01 association in four cohorts was performed with the GWAMA software tool^19^ and results were visualized with R software (3.3.2)^15^.

## RESULTS

### GENOME-WIDE ASSOCIATION ANALYSIS OF PENICILLIN ALLERGY

To discover genetic factors that may predispose to penicillin allergy, we conducted a genome-wide association study (GWAS) of 19.1 million single-nucleotide polymorphisms (SNPs) and insertions/deletions in UKBB and EstBB (minor allele frequency filter in both cohorts MAF > 0.1%). Cases were defined as participants with a Z88.0 ICD10 code (“Allergy status to penicillin”) for a reported history of penicillin allergy. In total, we identified 15,690 unrelated individuals (4.2% of the total cohort size of 377,545) in UKBB with this diagnostic code. However, the corresponding number of cases in EstBB was only 7 (0.02% of the total cohort size of 32,608) suggesting heterogeneity in the use of the Z88.0 ICD10 code in different countries. We therefore also identified participants that had self-reported drug allergy at recruitment in EstBB and categorized the EstBB self-reported reactions by drug class, using the Anatomical Therapeutic Chemical (ATC) Classification System code J01C* (beta-lactam antibacterials, penicillins) to match this to the respective Z88.0 ICD10 code. This resulted in 961 (2.9%) unrelated cases with penicillin allergy in EstBB. We validated the approach in EstBB by evaluating the association between the number of filled (i.e. prescribed and purchased) penicillin (using the ATC code J01C*) prescriptions per person and self-reported penicillin allergy. Using Poisson regression analysis, we identified a negative effect on the number of filled penicillin prescriptions among individuals with self-reported allergy in EstBB (P-value 2.41×10^-15^, Estimate −0.18 i.e. prescription count is 16% lower for individuals with penicillin allergy).

We then meta-analyzed the results of the GWASes in these two cohorts, weighing effect size estimates using the inverse of the corresponding standard errors. We identified a strong genome-wide significant (p < 5×10^-8^) signal for penicillin induced allergy (defined as ICD10 code Z88.0 or reported allergy to drugs in ATC J01C* class) on chromosome 6 in the major histocompatibility complex (MHC) region (lead variant rs114892859, MAF(EstBB) = 0.7%, MAF(UKBB) = 2%, P = 4.59×10^-29^, OR 1.59 95% CI 1.47-1.73) (**Figure 1; Table S1**).

**Figure 1.**
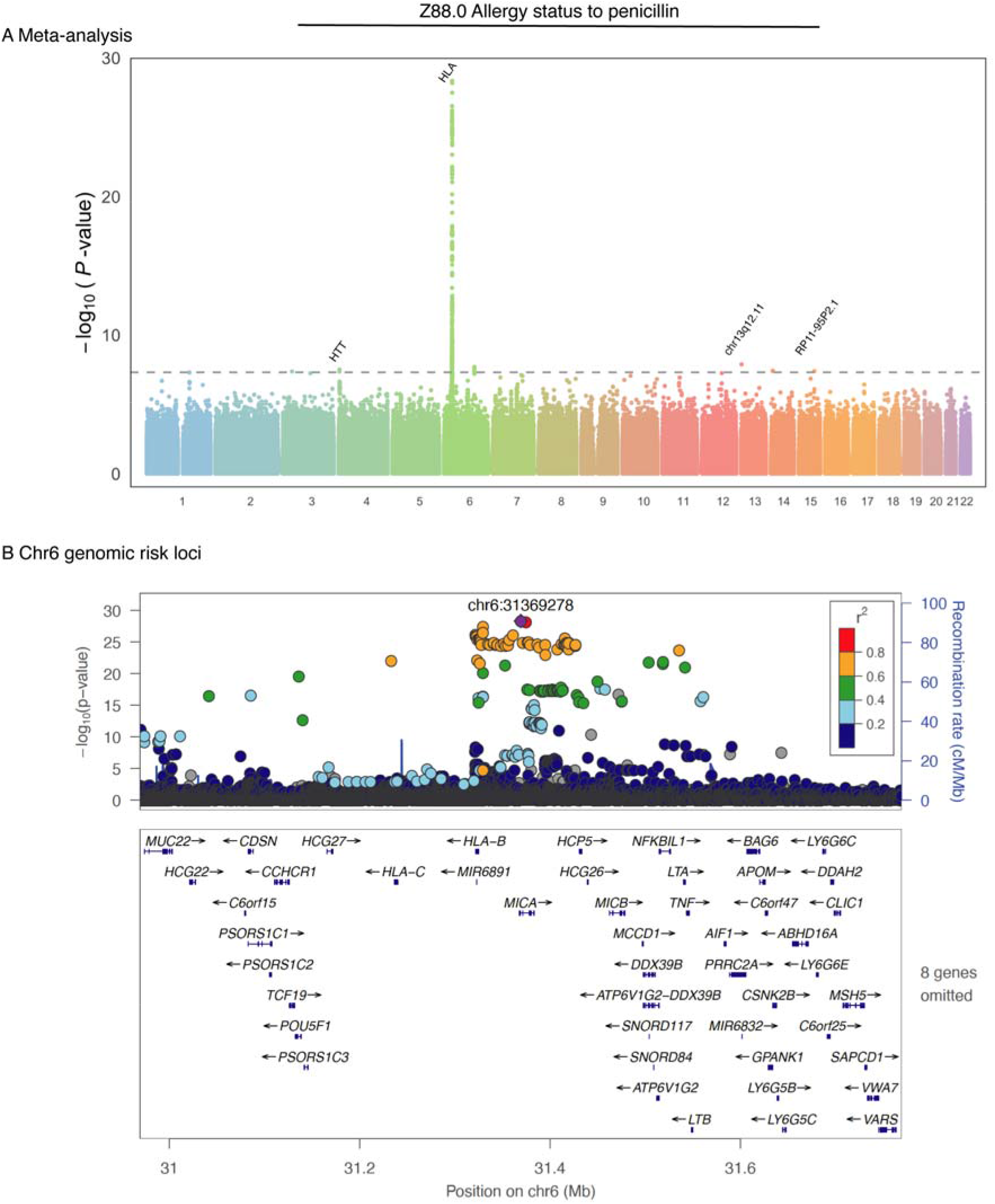
Manhattan plot (A) and HLA locus (B) of the genome-wide association study of allergy status to penicillin. The X-axes indicate chromosomal positions and Y-axes −log_10_ of the P-values **(A)** Each dot represents a single nucleotide polymorphism (SNP). The dotted line indicates the genome-wide significance (P-value<5.0×10^-8^) P-value threshold. **(B)** SNPs are colored according to their linkage disequilibrium (LD; based on the 1000 Genome phase3 EUR reference panel) with the lead SNP. The SNP marked with a purple diamond is the top lead SNP rs114892859 identified depending on LD structure.

### FINE-MAPPING THE PENICILLIN ALLERGY-ASSOCIATED HLA LOCUS

To further fine-map the causal variant of the identified association with penicillin allergy, we performed a functional annotation analysis with FUMA (Functional Mapping and Annotation of Genome-Wide Association Studies)^20^. We detected an independent intronic lead SNP for the penicillin allergy meta-analysis (GWAS lead variant rs114892859, P-value 2.21×10^-28^) in the *MICA* gene (**Figure 1, B**). When testing the SNP for expression quantitative trait locus (eQTL) associations in blood based on data from the eQTLGen Consortium^21^, the variant appeared to be associated with the expression levels of several nearby genes, with the most significant being *PSORS1C3* (P-value 8.10×10^-62^) and *MICA* (P-value 1.21×10^-52^) (**Table S2**). We further performed an *in silico* investigation of the lead SNP rs114892859 and its best proxy (rs144626001, the only proxy with r^2^>0.9 in UKBB and EstBB) in HaploReg v4 to explore annotations and impact of the non-coding variant^22^. In particular rs114892859 had several annotations indicative of a regulatory function, including its location in both promoter and enhancer marks in T-cells and evidence of RNA polymerase II binding^23,24^. Interestingly, its proxy is more likely to be deleterious based on the scaled Combined Annotation Dependent Depletion (CADD) score (scaled score of 15.78 for rs144626001 (C/T) and 4.472 for rs114892859 (G/T))^25,26^.

Due to the high LD in the MHC region, we used imputed SNP to HLA typing data available at four-digit resolution^27^ for up to 22,554 and 488,377 individuals from the Estonian and UK cohorts, respectively, to further fine-map the identified HLA association with penicillin allergy. In both cohorts a shared total of 103 alleles at four-digit level were present for all of the MHC class I genes (*HLA-A, HLA-B, HLA-C*) and 59 alleles for three of the classical MHC class II genes (*HLA-DRB1, HLA-DQA1, HLA-DQB1*). To assess the variation in the frequencies of the HLA alleles in different populations, we compared the obtained allele frequencies in both cohorts (**Table S3**) with the frequencies of HLA alleles in different European, Asian and African populations reported in the HLA frequency database (**Figure S2 and S3, Table S4**).

We then used an additive logistic regression model to test for associations between different four-digit HLA alleles and penicillin allergy in UKBB and EstBB. The results of both cohorts were meta-analyzed and P-values passing a Bonferroni correction (0.05/162 = 3.09×10^-4^, where 162 is the number of meta-analyzed HLA alleles) were considered significant (**Table S5**). One of the three results that surpassed the significance threshold had discordant effects in the two cohorts and one had a marginally significant association (P-value 2.81×10^-4^, **Table S5**). The strongest association we detected for penicillin allergy was the HLA-B*55:01 allele (P-value 4.63×10^-26^; OR 1.47 95% CI 1.37-1.58), which is tagged (r^2^>0.95) by the GWAS lead variant rs114892859 (**Table S6**).

### REPLICATION OF HLA-B*55:01 ASSOCIATION WITH PENICILLIN ALLERGY

To further confirm association with penicillin allergy we analyzed the association of the HLA-B*55:01 allele with self-reported penicillin allergy among 87,996 cases and 1,031,087 controls from the 23andMe research cohort. We observed a strong association (P-value 1.00×10^-47^; OR 1.30 95% CI 1.25-1.34; **Figure 2**) with a similar effect size as seen for the HLA-B*55:01 allele in the meta-analysis of the EstBB and UKBB. We obtained further confirmation for this association from the published dataset of Vanderbilt University’s biobank BioVU, where the HLA-B*55:01 allele was associated with allergy/adverse effect due to penicillin among 58 cases and 23,598 controls (P-value 1.79×10^-2^; OR 2.15 95% CI 1.19-6.5; **Figure 2**)^28^. Meta-analysis of results from discovery and replication cohorts demonstrated a strong association of the HLA-B*55:01 allele with penicillin allergy (P-value 2.23×10^-72^; OR 1.33 95% CI 1.29-1.37; **Figure 2**).

**Figure 2.**
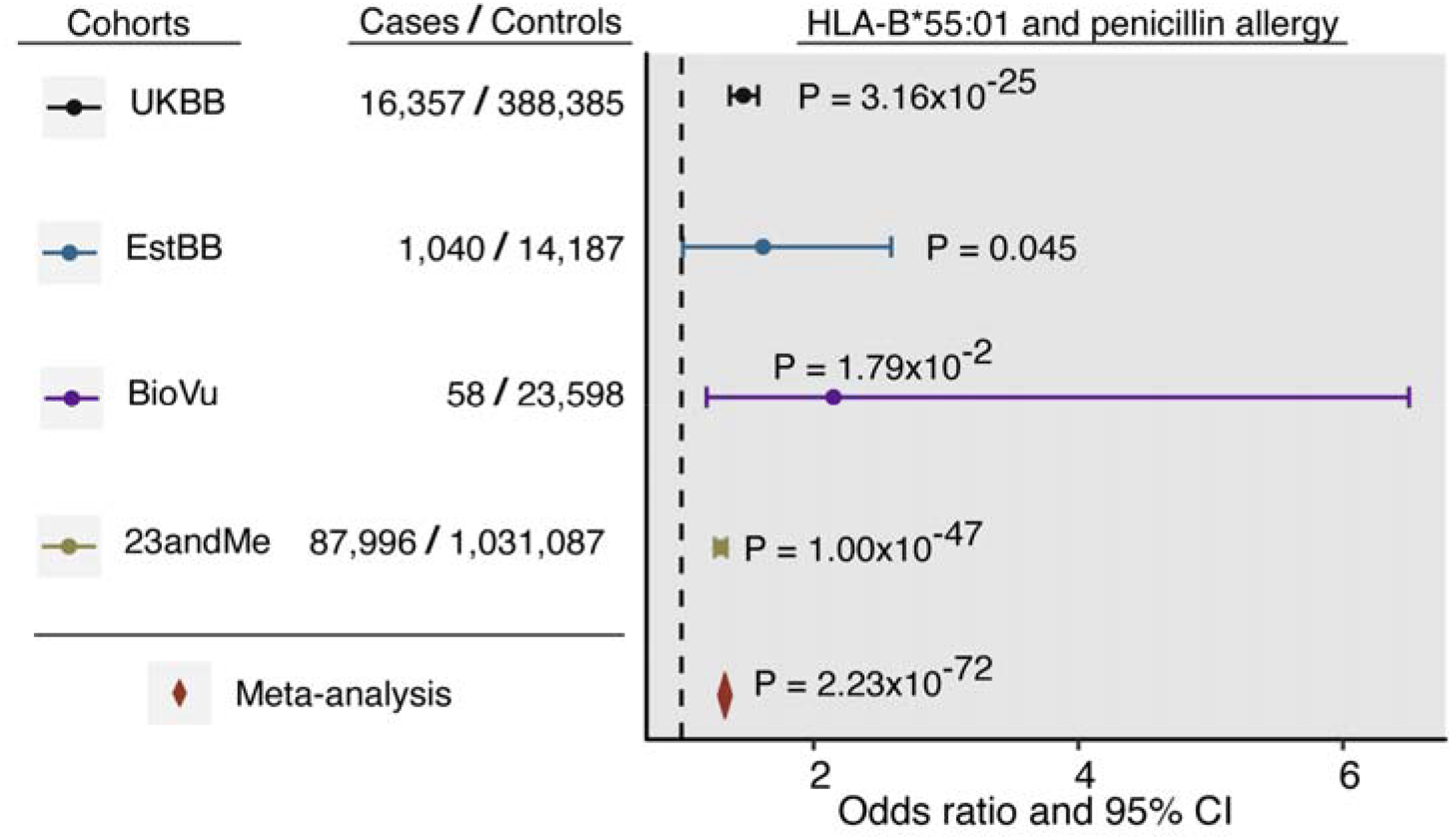
HLA-B*55:01 allele association with penicillin allergy-. The odds ratios (dots) and 95% confidence intervals (CI, horizontal lines) for HLA allele associated with penicillin allergy. The plot is annotated with P-values and case-control numbers. Color coding blue and black indicates the results for discovery cohorts Estonian UK biobank and replication results of the HLA*B-55:01 allele in 23andMe research cohort (green) and Vanderbilt University’s biobank BioVU (purple). Results of the meta-analysis of all four cohorts is indicated with a diamond (red).

### FURTHER ASSOCIATIONS AT HLA-B*55:01

Finally, we used the Open Targets Genetics platform’s UKBB PheWAS data^29^ to further characterize the association of the GWAS lead variant (and HLA-B*55:01 allele tag-SNP) rs114892859 (**Table S6**) with other traits. We found strong associations with lower lymphocyte counts (P-value 9.21×10^-14^, −0.098 cells per nanoliter, per allergy-increasing T allele) and lower white blood cell counts (P-value 3.17×10^-9^, −0.078 cells per nanoliter, per allergy-increasing T allele). To confirm this association, we extracted data on lymphocyte counts from the EHRs data of 4,567 EstBB participants (see **Supplementary methods**), and observed the same inverse association of the HLA-B*55:01 allele with lymphocyte counts (−0.148 cells per nanoliter, per T allele; P-value=0.047).

## DISCUSSION

In the present study, we identify a strong genome-wide significant association of the HLA-B*55:01 allele with penicillin allergy using data from four large cohorts: UKBB, EstBB, 23andMe and BioVu.

Hypersensitivity or allergic reactions to medications are type B adverse drug reactions that are known to be mediated by the immune system. One major driver of hypersensitivity reactions is thought to be the HLA system, which plays a role in inducing the immune response through T cell stimulation, and is encoded by the most polymorphic region in the human genome^30^. Genetic variation in the HLA region alters the shape of the peptide-binding pocket in HLA molecules, and enables their binding to a vast number of different peptides – a crucial step in the adaptive immune response^31^. However, this ability of HLA molecules to bind a wide variety of peptides may also facilitate binding of exogenous molecules such as drugs, potentially leading to off-target drug effects and immune-mediated ADRs^32^. The precise mechanism of most HLA-drug interactions remains unknown, but it seems that T cell activation is necessary for the majority of HLA-mediated ADRs^32–34^. Despite the increasing evidence for a role of the HLA system in drug-induced hypersensitivity, much is still unclear, including how genetic variation in the HLA region predisposes to specific drug reactions.

Penicillin is the most common cause of drug allergy, with clinical manifestations ranging from relatively benign cutaneous reactions to life-threatening systemic syndromes^7,8^ There is a previous GWAS on the immediate type of penicillin allergy, where a borderline genome-wide significant protective association of an allele of the MHC class II gene *HLA-DRA* was detected and further replicated in a different cohort^35^ Here we detect a robust association between penicillin allergy and an allele of the MHC class I gene *HLA-B*. The allele and its tag-SNP were also associated with lower lymphocyte levels and overlapped with T cell regulatory annotations, which suggests that the variant may predispose to a T-cell-mediated, delayed type of penicillin allergy. MHC I molecules are expressed by almost all cells and present peptides to cytotoxic CD8+ T cells, whereas MHC II molecules are expressed by antigen-presenting cells to present peptides to CD4+ T helper lymphocytes^31,34^ There are several examples of MHC I alleles associated with drug-induced hypersensitivity mediated by CD8+ T cells^34,36,37^ The involvement of T cells in delayed hypersensitivity reactions has been shown by isolating drug reactive T cell clones^38^, and cytotoxic CD8+ T cells have been shown to be relevant especially in allergic skin reactions^39–41^. More than twenty years ago, CD8+ T cells reactive to penicillin were isolated from patients with delayed type of hypersensitivity to penicillin^42^. The association with the HLA-B*55:01 allele detected in our study might be a relevant factor in this previously established connection with CD8+ T cells. The HLA-B*55:01 allele, together with other HLA-B alleles that share a common “E pocket sequence”, has previously been associated with increased risk for eosinophilia and systemic symptoms, Stevens-Johnson Syndrome and toxic epidermal necrolysis (SJS/TEN) among patients treated with nevirapine^43^. The underlying mechanism in penicillin allergy remains a question and various models have been proposed for T-cell-mediated hypersensitivity^36,41^. For example, the hapten model suggests that drugs may alter proteins and thereby induce an immune response^36,44^ – penicillins have been shown to bind proteins^44,45^ to form hapten–carrier complexes, which may in turn elicit a T cell response^46^ Drugs may also bind with MHC molecules directly. For example, abacavir has been shown to bind non-covalently to the peptide-binding groove of HLA-B*57:01, leading to a CD8+ T cell-mediated hypersensitivity response^47^.

It is being increasingly recognized that the involvement of HLA variation in hypersensitivity reactions goes beyond peptide specificity. Other factors, such as effects on HLA expression that influence the strength of the immune response have also been described^48^ The analysis of eQTLs based on the data of the eQTLGen Consortium^21^ revealed that the T allele of the lead SNP rs114892859 identified in our GWAS of penicillin allergy appears to be associated with the expression of several nearby genes, including lower expression of both *HLA-B* and *HLA-C*, and an even stronger effect on RNA levels of *PSORS1C3* and *MICA* (**Table S2**). Interestingly, variants in the *PSORS1C3* gene have been associated with the risk of allopurinol, carbamazepine and phenytoin induced SJS/TEN hypersensitivity reactions^49^. *MICA* encodes the protein MHC class I polypeptide-related sequence A^50^ which has been implicated in immune surveillance^51,52^. Our findings therefore support the observation that variants associated with expression of HLA genes may contribute to the development of hypersensitivity reactions. We detect strong evidence for the involvement of HLA-B*55:01 in penicillin allergy, and a marginally significant association in the MHC II gene DRB1, although both need further functional investigation to explore their exact roles and mechanisms in the induced response.

The main limitation of this study is the unverified nature of the phenotypes extracted from EHRs and self-reported data in the biobanks. Previous work has found that most individuals labeled as having beta-lactam hypersensitivity may not actually have true hypersensitivity^7,8,53^ Nevertheless, despite the possibility that some cases in our study may be misclassified, we detect a robust HLA association that was replicated in several independent cohorts against related phenotypes. The increased power arising from biobank-scale sample sizes therefore mitigates some of the challenges associated with EHR data. The robustness of the genetic signal across cohorts with orthogonal phenotyping methods, ranging from EHR-sourced in UKBB to various forms of self-reported data in EstBB and 23andMe, also supports a true association. Finally, the modest effect size of the HLA-B*55:01 allele (OR 1.33), particularly when compared to effect sizes of HLA alleles with established pharmacogenetic relevance^54–56^, suggests that this variant in isolation is unlikely to have clinically meaningful predictive value. However, further phenotypic refinement, including investigation of specific penicillin-based medicines and specific types of drug reactions, may yield more clinically actionable insight. Our work also provides the foundation for further studies to investigate the application of a polygenic risk score^57^ (which combines the effects of many thousands of trait-associated variants into a single score), possibly in combination with phenotypic risk factors, in identifying individuals at elevated risk of penicillin allergy.

In summary, our results provide novel evidence of a robust genome-wide significant association of HLA and the HLA-B*55:01 allele with penicillin allergy. Further phenotypic refinement, including investigation of specific penicillin-based medicines and specific types of drug reactions, may also yield more clinically actionable insight.

## Acknowledgements

This study has been supported by grants from the Estonian Research Council grant numbers PRG184, PRG687, IUT20-60 and IUT24-6, and the Oak Foundation. This work was carried out in part in the High Performance Computing Center of University of Tartu. We acknowledge the Finnish SISu Project and principal investigators Aarno Palotie, Jaana Suvisaari, Veikko Salomaa, and Priit Palta for sharing the Finnish imputation reference panel. This research has been conducted using the UK Biobank Resource under Application Number 11867. We thank the research participants of 23andMe for their contribution to this study and the 23andMe Research Team. We further thank all the participants and staff of the Estonian, UK and Vanderbilt university biobanks for their contribution to this research. J.B. is supported by funding from the Rhodes Trust, Clarendon Fund and the Medical Sciences Doctoral Training Centre, University of Oxford. J.C.C. is funded by the Oxford Medical Research Council Doctoral Training Partnership (Oxford MRC DTP) and the Nuffield Department of Clinical Medicine, University of Oxford. C.M.L. is supported by the Li Ka Shing Foundation; WT-SSI/John Fell funds; the NIHR Biomedical Research Centre, Oxford; Widenlife; and NIH (5P50HD028138-27). M.V.H. works in a unit that receives funding from the MRC and is supported by a British Heart Foundation Intermediate Clinical Research Fellowship (FS/18/23/33512) and the National Institute for Health Research Oxford Biomedical Research Centre. Computation used the Oxford Biomedical Research Computing (BMRC) facility, a joint development between the Wellcome Centre for Human Genetics and the Big Data Institute supported by Health Data Research UK and the NIHR Oxford Biomedical Research Centre. Financial support was provided by the Wellcome Trust Core Award Grant Number 203141/Z/16/Z. The views expressed are those of the author(s) and not necessarily those of the NHS, the NIHR or the Department of Health.

## Author Contributions

K.K., L.M. and J.F. designed the study. R.M., M.L., Y.L., S.R., A.M. and T.E. supervised and generated genotype data or HLA typing data. D.S. and S.L. generated allergy data from free-text. K.K., J.B., M.L., T.J., J.C.C., J.F, W.W., A.A., performed the data analysis. K.K., J.B., M.V.H. C.M.L., R.M., L.M., J.C.C. and J.F. conducted data interpretation. K.K. prepared the figures and tables. K.K, J.B., L.M. and J.F. drafted the manuscript. K.K., J.B., M.V.H. C.M.L., M.L., R.M., L.M., J.C.C., W.W., A.A. and J.F. reviewed and edited the manuscript. All authors contributed to critical revisions and approved the final manuscript.

The following members of the 23andMe Research Team contributed to this study: Michelle Agee, Stella Aslibekyan, Robert K. Bell, Katarzyna Bryc, Sarah K. Clark, Sarah L. Elson, Kipper Fletez-Brant, Pierre Fontanillas, Nicholas A. Furlotte, Pooja M. Gandhi, Karl Heilbron, Barry Hicks, David A. Hinds, Karen E. Huber, Ethan M. Jewett, Yunxuan Jiang, Aaron Kleinman, Keng-Han Lin, Nadia K. Litterman, Marie K. Luff, Jennifer C. McCreight, Matthew H. McIntyre, Kimberly F. McManus, Joanna L. Mountain, Sahar V. Mozaffari, Priyanka Nandakumar, Elizabeth S. Noblin, Carrie A.M. Northover, Jared O’Connell, Aaron A. Petrakovitz, Steven J. Pitts, G. David Poznik, J. Fah Sathirapongsasuti, Anjali J. Shastri, Janie F. Shelton, Suyash Shringarpure, Chao Tian, Joyce Y. Tung, Robert J. Tunney, Vladimir Vacic, Xin Wang, Amir S. Zare.

